## Supplemental File 1 for "Rapid, high-throughput phenotypic profiling of endosymbiotic dinoflagellates (Symbiodiniaceae) using benchtop flow cytometry"

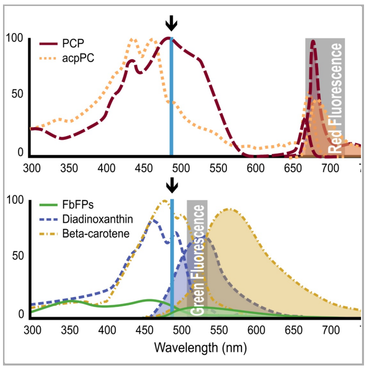


CV5CW82W

### Rapid, high-throughput phenotypic profiling of endosymbiotic dinoflagellates (Symbiodiniaceae) using benchtop flow cytometry V.(cv5cw82w)

Bastian
Colin J Anthony[1], Colin Lock[1], Bentlage[1]

[1]Marine Laboratory, University of Guam, Mangilao, Guam, USA

Colin J Anthony
Marine Laboratory, University of Guam

**Protocol Info:** Colin J Anthony, Colin Lock, Bastian Bentlage . Rapid, high-throughput phenotypic profiling of endosymbiotic dinoflagellates (Symbiodiniaceae) using benchtop flow cytometry. **protocols.io** https://protocols.io/view/rapid-high-throughput-phenotypic-profiling-of-endo-cv5cw82w

**MANUSCRIPT CITATION:**
Anthony CJ, Lock C, Bentlage B. Rapid, high-throughput phenotypic profiling of endosymbiotic dinoflagellates (Symbiodiniaceae) using benchtop flow cytometry. PLOS ONE. In review.

**Created:** Jun 22, 2023

**Last Modified:** Jun 22, 2023

**PROTOCOL integer ID:** 83844

**Keywords:** flow cytometry, autofluorescence, photopigment, dinoflagellate, Symbiodiniaceae, coral, Cnidaria, symbiosis, Symbiodinium, fluorescence

#### ABSTRACT

Endosymbiotic dinoflagellates (Family Symbiodiniaceae) are the primary producer of energy for many cnidarians, including corals. The intricate coral-dinoflagellate symbiotic relationship is becoming increasingly important under climate change, as its breakdown leads to mass coral bleaching and often mortality. Despite methodological progress, assessing the phenotypic traits of Symbiodiniaceae *in-hospite* remains a complex task. Bio-optics, biochemistry, or “-omics” techniques are expensive, often inaccessible to investigators, or lack the resolution required to understand single-cell phenotypic states within endosymbiotic dinoflagellate assemblages. To help address this issue, we developed a protocol that collects information on cell autofluorescence, shape, and size to simultaneously generate phenotypic profiles for thousands of Symbiodiniaceae cells, thus revealing phenotypic variance of the Symbiodiniaceae assemblage to the resolution of single cells. As flow cytometry is adopted as a robust and efficient method for cell counting, integration of our protocol into existing workflows allows researchers to acquire a new level of resolution for studies examining the acclimation and adaptation strategies of Symbiodiniaceae assemblages.

#### IMAGE ATTRIBUTION

Colin Anthony 2023

#### GUIDELINES

We focus on the characterization of Symbiodiniaceae associated with reef-building corals (Scleractinia); however, we have had success using this for upside-down jellyfish (*Cassiopea*). Therefore, with slight modifications this protocol may be used for flash-frozen and live cells for both calcifying and gelatinous cnidarians.
Each instrument, environment, and organism is different, so it is important to optimize this protocol locally.
Samples are prone to degradation, so it is important to work efficiently and consistently.
It is vital the same equipment is used in the same environment each time to avoid batch effect.
If using in a large multi-factorial project, mixed batches are recommended to avoid batch effect.
This protocol pairs well with cell counting methods, ITS2 metabarcoding, photopigment spectrophotometry, and coral morphometric methods, so we would suggest integrating this into more complex data structures as necessary.
Krediet et al. (2015) and Apprill et al. (2007) were pivotal in providing a framework for developing this protocol.

##### CITATION

Krediet CJ, DeNofrio JC, Caruso C, Burriesci MS, Cella K, Pringle JR (2015). Rapid, Precise, and Accurate Counts of Symbiodinium Cells Using the Guava Flow Cytometer, and a Comparison to Other Methods.. PloS one.

LINK

https://doi.org/10.1371/journal.pone.0135725

##### CITATION

Apprill AM, Bidigare RR, Gates RD (2007). Visibly healthy corals exhibit variable pigment concentrations and symbiont phenotypes. Coral Reefs.

LINK

https://doi.org/10.1007/s00338-007-0209-y

#### MATERIALS

**Preservation**

Instruments:

- Ultra low freezer -80 °C (TSXSeries powered by V-Drive) *

Equipment:

- Cryogenic storage Dewar flask(Thermo Scientific Nalgene 4150-2000 All-plastic Dewar flask, 2 L) *
- Cryogenic storage Liquid Dewar (VWR® CryoPro® Liquid Dewar, L Series, 3.8 L) *
- Wire cutters **
- Sample bag (Whirl-Pak® Write-On Bags - 4 oz. - Yellow Tape) **

- Thermo Scientific screw cap micro tubes
  Tether screw cap w O-ring, natural (3466NKS)
  1.5 mL screw cap tube, NonKnurl, NonSkirted, Natural (3466NKS)

Chemicals:

- Liquid nitrogen (LN2)

**Sample Prep**

Instruments:

- Precision balance (METTLER TOLEDO PB303-S)
- Benchtop shaking incubator (222DS)
- Air Compressor (TC-20T) **
- Airbrush (TJ-180 ) **
- Ice maker
- Weigh boats
- Bead beater (MiniBeadBeater Plus)
- Centrifuge (Eppendorf 5425 R)
- Test tube shaker (Lab Dancer S000)

Equipment:

- 50 mL falcon tubes **
- Dissecting forceps **
- Small cooler (Rubbermaid 2A21)
- 50 mL tube storage rack (4- Sides plastic Micro Tube Rack)
- 1.5 mL tube storage rack (80-Place Lab Storage Rack LC537)
- 10 mL 20G1 latex free syringe with PrecisionGlide needle
- Thermo Scientific screw cap micro tubes
  Tether screw cap w O-ring, natural (3466NKS)
  1.5 mL screw cap tube, NonKnurl, NonSkirted, Natural (3466NKS)
- 96 round bottom microwell plate (Nunc™ 96-Well Polystyrene Round Bottom Microwell Plates 262162)
- 100-1000 uL pipette (BioPette™ Plus P3942-1000)
- 20-200 uL pipette (BioPette™ Plus P3942-200)
- 2-20 uL pipette (BioPette™ Plus P3942-20)
- 20-200 uL universal pipet tips (VWR® 76322-150)
- 100-1000 uL universal pipet tips (VWR® 76322-154)

Chemicals:

- Crushed ice
- Filtered Seawater (FSW)
- Deionized water (DI)
- Lauryl sulfate (SDS: Sodium dodecyl sulfate) sodium salt (L 4390)

**Cytometry**

Instruments:

- **Flow cytometer (Luminex Guava easyCyte 6HT-2L)**
- Computer (hp with Intel Core i7 processer)

Equipment:

- 20-200 uL pipette (BioPette™ Plus P3942-200)
- 2-20 uL pipette (BioPette™ Plus P3942-20)
- 20-200 uL universal pipet tips (VWR® 76322-150)

- CountBright™ Plus Absolute Counting Beads (C36995)
- Guava® easyCheck™ Kit (4500-0025) (https://www.luminexcorp.com/guava-easycheck-kit/?wpdmdl=40649)

Chemicals:

- Deionized water (DI)
- 100% bleach
- Guava Instrument Cleaning Fluid (30-00133) (ICF)

Programs:

- guavaSoft v4.0
  - Guava Clean v3.4
  - InCyte v4.0

**Post-processing**

Instruments:

- Computer (hp with Intel Core i7 processer)

Programs:

- guavaSoft v4.0
  - Guava Clean v3.4
  - InCyte v4.0
- R v4.1.2 (R Core Team 2021)
  - dplyr v1.0.10 (Wickham et al. 2022)
  - tidyr v1.2.0 (Wickham and Girlich 2022)
  - readr v2.1.2 (Wickham et al. 2022)
  - ggplot2 (Wickham 2016)
  - ggpubr v0.4.0 (Kassambara 2020)
  - cowplot v1.1.1 (Wilke 2020)
- RStudio v1.3.1073 (RStudio Team 2020)

** Unnecessary if working with live cells*

*** For cnidarians with calcium carbonate skeleton*

#### SAFETY WARNINGS

This method is not particularly dangerous, though we do recommend using a mask to avoid inhaling any foreign particles during airbrushing and using extreme caution when handling cutters, needles, and liquid nitrogen.

BEFORE START INSTRUCTIONS

**Ensure filtered seawater (FSW) is fresh**

**Heat SDS solution** with benchtop shaking incubator to resuspend salt

180 rpm, 70°C, 00:10:00

- 50 mL SDS solution = 7 mL FSW + 43 mL DI + 0.04 g SDS

**Clean the flow cytometer** before and after each use

- GuavaSoft 4.0 uses Guava Clean 3.4, which walks you through cleaning process
- Two pre-cleans is helpful and if the machine sat over the weekend, cleaning the capillary is also necessary
- Throw out waste bottle and make sure other bottle has at least ⅔ ICS

**Verify proper instrument calibration and gain settings with easyCheck and CountBright fluorescent beads**

- Once the protocol has been implemented locally, use fluorescent beads to verify that fluorescent readings are consistent. Set up a worklist with bins that show the expected location of each fluorescent bead. This file is used to periodically check cytometry calibration.
- Periodically before collecting data (every 1-3 months), fill a single well with 1uL of fluorescent beads. Run the cytometer, and while it is running, verify that fluorescent beads are within the expected bin, if not, adjust the gain settings to put fluorescent readings within your predefined gates.
- Reagents and instructions come with easyCheck kit.

Note

WARNING: If runs are far apart, and not verified for proper calibration with CountBright fluorescent beads, data between runs cannot be compared. Data between multiply cytometry runs MUST be verified for proper calibration with fluorescent beads.

**Clean air brush** and make sure needle is still present*

**Chill centrifuge** 5000 rcf, 0°C

**Fill cooler with ice**

**Locate samples and label all tubes** (50 mL falcon tubes* and 1.5 mL screw top tubes)

#### Sample Preservation

3m

1 **Sample tissue from cnidarian**

1m

- If calcifying coral, we suggest 3 pieces at around 2 $cm^3$ sampled with wire cutters or hammer and chisel
- If gelatinous cnidarian, 0.05 - 0.1 g of tissue sampled with sterile scissors works well

Note

If desired, cut an additional piece from the sample for DNA extraction or other relevant methodologies and workflows.

Protocol

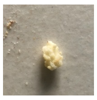

NAME

**Coral DNA Extraction - Modified DNeasy PowerSoil Pro Kit**

CREATED BY

Luigi Colin

PREVIEW

2 **Store tissue in bag or tube** 1m

- For calcifiers, Whirl-Pak sample bags work well
- For gelatinous cnidarians, 1.5 mL screw top tubes are more effective (this allows for immediate processing when ready)

3 **Flash-freeze tissue** in Cryogenic storage Dewar flask with LN2 -320 °C 1m

- Store samples in ultra low freezer -80 °C for extended preservation

#### Sample Preparation 2h

4 **Before Start:** 30m

Ensure filtered seawater (FSW) is fresh (~1 L of filtered seawater is sufficient)

Heat SDS solution with benchtop shaking incubator to resuspend salt

180 rpm, 70°C, 00:10:00

- 50 mL SDS solution = 7 mL FSW + 43 mL DI + 0.04 g SDS
- Make sure solution cools to ambient temperature before use

Clean the flow cytometer before and after each use

- GuavaSoft 4.0 uses Guava Clean 3.4, which walks you through cleaning process
- Two pre-cleans is helpful and if the machine sat over the weekend, cleaning the capillary is also necessary
- Throw out waste in waste bottle and make sure other bottle has at least ⅔ ICS

easyCheck may also be necessary if machine has sat for more than a week

- reagents and instructions come with easyCheck kit

Verify proper instrument calibration and gain settings with CountBright fluorescent beads

- Once the protocol has been implemented locally, use fluorescent beads to verify that fluorescent readings are consistent. Set up a worklist with preset bins that show the expected location of each fluorescent bead. (This is the same binning process used to bin Symbiodiniaceae cellular populations)

- Periodically before collecting data (every 1-3 months), fill a single well with 1uL of fluorescent beads. Run the cytometer, and while it is running, verify that the fluorescent beads are within the expected bin, if not, adjust the gain settings.

Clean air brush and make sure needle is still present*

Chill centrifuge 5000 rcf, 0°C

Fill cooler with ice

Locate samples and label all tubes (50 mL falcon tubes* and 1.5 mL screw top tubes)

- For 12 samples, you will need 12 falcon tubes, 12 1.5 mL screw top tubes, and 1 microwell plate
- Label one falcon tube and one 1.5 mL tube (If desired, add more 1.5 mL tubes as added technical replicates) for each sample being processed

5 **Prepare tissue slurry (calcifying cnidarians only)** 1h

**Before start:**

- Make sure spray gun compressor is turned on
- Have somewhere to store skeletal fragments after air-brushing (e.g. small weigh boats)
- Be prepared to work efficiently as samples degrade quickly once removed from the freezer

5.1 Locate samples in ultra-low freezer and remove up to 3 samples at a time

5.2 Cut a ~1-2 cm piece from coral fragment (this will vary slightly depending on species)

5.3 Use forceps to hold coral fragment just inside the mouth of a 50 mL falcon tube, then use an airbrush loaded with filtered seawater (FSW) to remove all coral tissue from the skeleton making sure to capture the tissue in the falcon tube.

- Depending on the species, this can take 5-20 mL of FSW
- Store falcon tube on ice in dim ambient lighting
- Place the remaining skeleton on a weigh boat (Keep track of it. You will need it to normalize cell densities at the end)

5.4 Repeat steps 5.1-5.3 with all samples

- Move quickly and take no more than 1 hour for all samples combined

5.5 Once all samples have been airbrushed, vortex and needle shear each tissue slurry until homogenized

- Ensure full homogeny of slurry. Allowing any settling or heterogeneity can skew data consistency: There should be no mucus clumps or visible chunks; however, it is normal for small skeletal fragments to settle at the bottom

5.6 Once a slurry has been homogenized, transfer 1 mL to a 1.5 mL screw top tube.

6 **Wash tissue slurries to be loaded in flow cytometer**

- If not using a calcifying cnidarian, add 1 mL of FSW to gelatinous tissue sample that was stored in a 1.5 mL screw-top tube

6.1 Bead beat 1.5 mL tubes 00:00:04 4s

6.2 Centrifugate samples 5000 rcf, 0°C, 00:04:00 4m

6.3 Remove 1 mL supernatant using 1000 mL pipette

- Do not pour supernatant out. Pellets are often loose and liquid does not empty completely

6.4 Resuspend pellets via repeated pipetting in 1 mL of FSW

6.5 Bead beat tubes again 00:00:04 4s

6.6 Centrifugate samples 5000 rcf, 0°C, 00:03:00 3m

6.7 Remove 1 mL supernatant using pipette

- Do not pour supernatant out. Pellets are often loose and liquid does not empty completely
- If you accidentally resuspend a pellet, centrifugate the sample for 30 seconds and try again.

6.8 Resuspend pellets via repeated pipetting in 1 mL of FSW and set samples to the side

#### Load the Cytometer

4s

7 **Prepare 96-microwell plate for cytometry**
**Before start , check for supplies:**

- 50 mL sufficient saltwater-freshwater solution (SFS)
  *25 mL FSW + 25 mL DI H2O = 50 ml SFS*
- 200 uL pipette tips
- 200 uL pipette
- 20 uL pipette
- Syringe with needle for needle sheering
- Vortexer

7.1 Load wells of microwell plate with 180 µL SFS.

- We describe this protocol with a 10x sample dilution (9 units SFS to 1 unit Sample). This volume of SFS will change as you optimize your dilutions
- We also do not recommend loading more than half a plate as fluorescent properties change after an extended period within the machine

> Note
>
> For us, a 10x dilution works well for a starting cell concentration of 100,000 - 200,000 cells/mL. It is also important to note that fluorescent signatures degrade within the machine, and cell counts become less reliable beyond row 4 (48 wells). Local protocol optimization is recommended, though we have had no issues with this protocol for 4 coral genera (*Acropora, Pavona, Pocillopora,* and *Porites*) and 1 jellyfish genus (*Cassiopea*).

7.2 Bead beat 00:00:04 a washed tissue sample then immediately load 20 µL of tissue homogenate into two wells preloaded with SFS before particulate settles 4s

- If there are any visible clumps left over from the symbiont pellet, use a combination of needle sheering and vortexing properly homogenize the sample; however, we do not recommend bead beating samples again as this risks lysing the algal cells
- Do not allow the samples to sit for any period of time in between mixing and loading. Any settling can skew data

7.3 Repeat step 7.2 until all samples have been loaded.

- We typically process 12-24 samples at a time. We do not recommend processing more than 24 samples in one run.

8 **Prepare worklist and set cytometry run settings for Guava Flow Cytometer**

> Note
>
> This step may be completed alongside or before step 4 to avoid the risk of sample degradation

8.1 In guavaSoft v4.0, open InCyte v4.0

8.2 Click "Edit Worklist"

8.3 Select the wells with loaded samples and click "Acquire Samples".

8.4 Set each wells setting to acquire for 180s with a maximum gated cell count of 2500.

> Note
>
> The gated cell count is the number of observations per sample that will be quantified within the R1 gate before moving on to the next sample. This R1 gate identifies symbiont cells based on Red off Blue fluorescence and side scatter based on our previous experiments, but this gate may need to be optimized for your symbiont community (See step 13).

8.5 Also set each well to have 2 technical replicates and 7 seconds of high energy stirring.

- This increases replication and prevents settling within the flow cytometer

8.6 Name each well

8.7 Click "Run Worklist"

9 **Load in appropriate method, settings, and compensation files**

**Starter Files**

Method:

Method.gsy

Settings & Compensation (Same file for both):

AcroSettingsCompensation.fcs

Note

Cytometer gain settings may need to be adjusted based on specific model of cytometer or target organism. Gain settings provided in this compensation file worked well for all samples. If gain is adjusted, save into Settings and Compensation file so gain is identical in all runs, which will prevent batch effects.

10 Click "Acquire"
*(If you would like to verify the integrity of your samples before starting the worklist, click "Adjust Settings")*

10.1 Follow the plate loading prompt

- Load DI H2O, ICF, and Bleach into the appropriate positions
- Place plate in the Guava Flow Cytometer in the appropriate orientation, as indicated by the marks in the loading tray

10.2 If you clicked "Adjust Settings"

1. Select the well of interest,
2. Verify your cells are in the appropriate region [Image Attached]
3. Once verified, click "Next Step"
4. click "Resume Worklist"

11 **Acquire samples** 5h

- A 48-well run should take ~ 05:00:00

Note

It is okay to leave it running at the end of the day, but make sure to pull out the tray and clean machine the next morning

Expected result

Cells of interest should fall in the upper right hand corner of the FSC-HLog to RED-B-HLog. These are your Symbiodiniaceae cells that were counted in the bin and are the source of the physiological signatures we are targeting. (This initial bin is broad, and a dataset-specific bin will be created post-hoc. See Step 13)

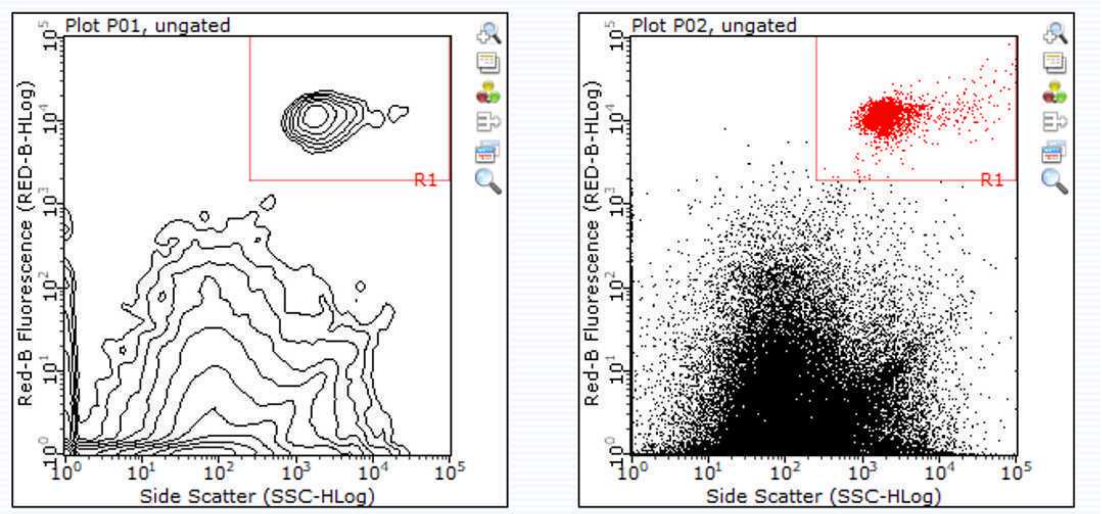


Two example plots illustrating a cluster of counted Symbiodiniaceae binned by the R1 gate.

#### Post-run Processing

##### 12 Export single-cell observations to determine dataset-specific binning threshold

12.1 When a run has finished, click the "Analyse" tab inside of InCyte.

12.2 In most cases, recent runs will be preloaded in "Analysed Data"; however, if your dataset of interest is missing, you may load your dataset in by clicking the blue folder that says "Open Analysed Group".

- Raw files are exported as YEAR-MONTH-DAY_at_HOUR-MINUTE-SECONDpm.fcs (e.g. 2022-10-04_01-38-41pm.fcs)
- Make sure the correct Method is applied to the analyzed data (Starter Method File: Method.gsy)

12.3 Highlight all wells of interest

- Click on one well, then click again and drag your mouse across your desired selection

12.4 Right click your highlighted selection and select "Export List Mode Data"

- Sometimes an error pops up saying that the file name has already been written. Don't worry, your files were successfully exported.

12.5 Locate your exported files of interest

- If combining multiple cytometry runs, we recommend placing all .csvs in the same file.

#### 13 Open RStudio to determine the dataset-specific symbiont binning threshold

Starter R Script is available here:

SetNewBinningThreshold.R

Example (Reduced Wells) List Mode Dataset:

Exp1_2022-09-21_at_11-08-48am.zip

Note

This can be computationally intensive for your computer, so if unable to complete this step as written, exporting a subset of wells to determine a binning threshold is typically fine. Statistical summaries would then be exported with the new bin, which is much less data-heavy.

13.1 Install and load in R Packages:

- dplyr v1.0.10 (Wickham et al. 2022)
- tidyr v1.2.0 (Wickham and Girlich 2022)
- readr v2.1.2 (Wickham et al. 2022)
- ggplot2 (Wickham 2016)
- ggpubr v0.4.0 (Kassambara 2020)
- cowplot v1.1.1 (Wilke 2020)

13.2 Import and combine all list mode data

Command

**Code written for R v4.1.2 (R Core Team 2021) in RStudio v1.3.1073 (RStudio Team 2020).**

```
(files <- fs::dir_ls("Directory/Exp1_2022-09-21_at_11-08-48am/",
glob="*.CSV"))
df <- read_csv(files, id="path")
head(df)
```

13.3 Replace "-" with "." and make our fluorescent signature of interest (red fluorescence off of the blue laser) into a numeric

Command

**Code written for R v4.1.2 (R Core Team 2021) in RStudio v1.3.1073 (RStudio Team 2020).**

```
names(df) <- gsub("-", ".", names(df), fixed=TRUE)


df$RED.B.HLog <- as.numeric(df$RED.B.HLog)
```

13.4 Plot the density of observations based on their relative red fluorescent intensity off of the blue laser

Command

**Code written for R v4.1.2 (R Core Team 2021) in RStudio v1.3.1073 (RStudio Team 2020).**

```
ggdensity(df, x = "RED.B.HLog", fill = "lightgray", rug = TRUE)+
  scale_x_continuous(limits = c(1.5, 5))
```

13.5 Determine your dataset-specific binning threshold to separate cells from other particles by identifying where x equals the minimum number of observations

Command

**Code written for R v4.1.2 (R Core Team 2021) in RStudio v1.3.1073 (RStudio Team 2020).**

```
DensityX < 4 & DensityX > 2
MinYDensity<- min(DensityY[DensityX < 4 & DensityX > 2])
MinYDensity
#0.003750236
which(DensityY == MinYDensity)
#334
DensityX[334]

#Visualize your threshold here
ggdensity(df, x = "RED.B.HLog", fill = "lightgray", rug =
TRUE)+
  scale_x_continuous(limits = c(1.5, 5))+
  geom_vline(xintercept = density(df$RED.B.HLog)$x[334])

#X Minimum = 3.005898
```

Expected result

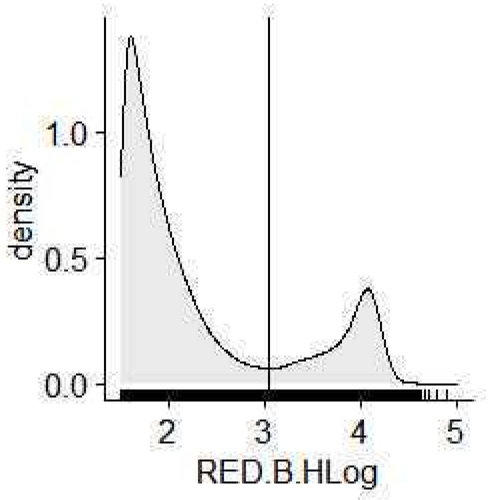


A vertical line should now illustrate your fluorescent threshold.

13.6 If desired, remove observations that do not fall within this binning threshold to have dataset, with every fluorescent profile for each fluorescent signature detected

- Metadata can be applied to this dataset based on file names

Command

**Code written for R v4.1.2 (R Core Team 2021) in RStudio v1.3.1073 (RStudio Team 2020).**

```
dfsym <- subset(df, RED.B.HLog>=3.005898)

ggdensity(dfsym, x = "RED.B.HLog", fill = "lightgray", rug =
TRUE)+
  scale_x_continuous(limits = c(1.5, 5))+
  geom_vline(xintercept = density(df$RED.B.HLog)$x[378])
```

Expected result

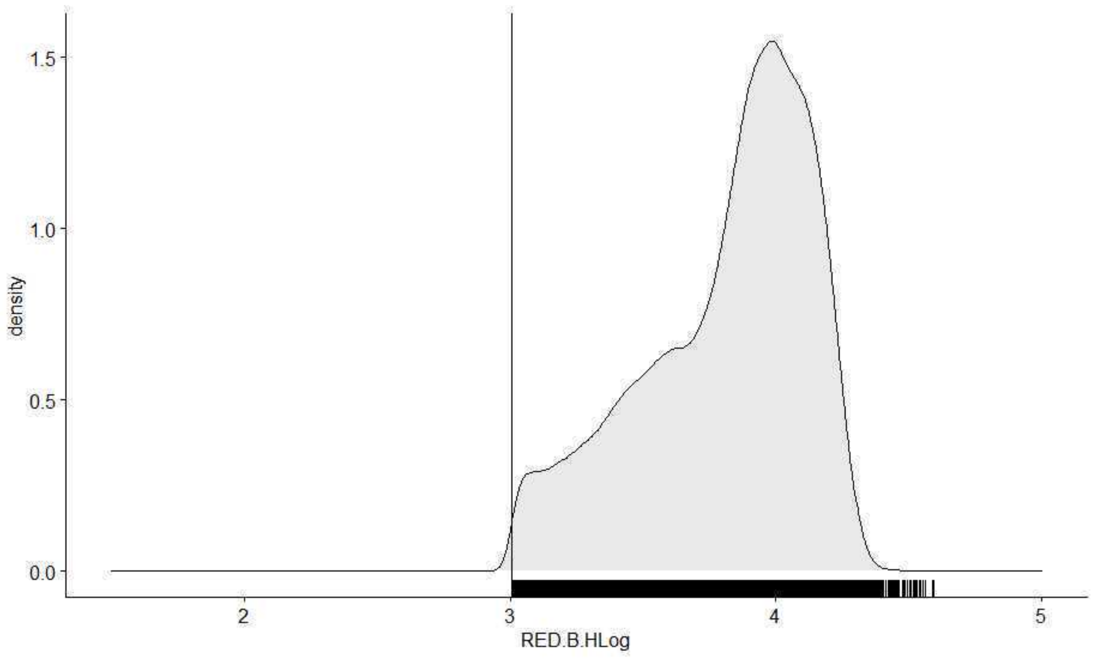


Density plot should no longer have any observations below the defined threshold.

13.7 Export filtered data to avoid the need for reprocessing. This data now contains all desired phenotypic data (red fluorescence, green fluorescence, forward scatter, side scatter).

Command

**Code written for R v4.1.2 (R Core Team 2021) in RStudio v1.3.1073 (RStudio Team 2020).**

```
write.csv(dat4sym,"IntendedDirectory/SubsetDataset.csv", row.names = FALSE)
```

**14** Using the threshold determined in step 13.5, manually adjust the bin titled "Symbiont" on the RED-B Fluorescence Histogram in InCyte:Analyse.

Note

Unfortunately there is no way to define a bin with numerical values in InCyte 4.0, so this binning is up to your best estimation. This is why we opt for a broad bin. This is also why it is important to apply the exact same method.gsy file across all analyzed groups to avoid creating batch effects.

If using multiple cytometry runs in your research, save your method with the correct "Symbiont" bin. You can apply this method to all analyzed .fcs files by clicking and dragging the method to the appropriate file within the InCyte:Analyse interface.

It is best to use the below bin for cell density calculations, and the previously filtered, calculated bin for fluorescent signatures.
More intimate code and robust datasets is available on:
https://github.com/AnthonyCuog/CytometryProtocol

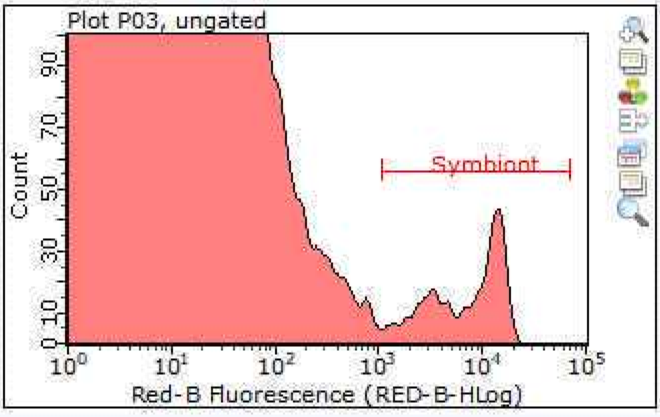


A resized "Symbiont" bin now sits at the estimated threshold for a random well.

15 Once the appropriate "Symbiont" bin has been applied to a dataset export a Group Stats .csv file

Note

The method file we have supplied in this protocol export the Cell Count, % Observation Included in Bin, Cellular Concentration, RED-B-HLog Mean, RED-B-HLog Median, RED-B-HLog %CV, GRN-B-HLog Mean, GRN-B-HLog Median, and GRN-B-HLog %CV.

15.1 On the left side of InCyte, click "Show Group Stats"

15.2 Click "Setup"

15.3 Remove the checkmarks for each empty field

15.4 Click "Done"

15.5 Click "Export to .csv" and save in desired location

15.6 The fluorescence readings are now ready to be used! Label your numbers appropriately, combine with other files, and apply any necessary metadata

#### Concentration Normalization

15.7 To determine the cell density, multiply the number exported (Concentration) in step 15 by your dilution and slurry volume, and then normalize your concentration to a surface area for calcifying Cnidaria (e.g. Koch et al. 2021) or protein content for non-calcifying Cnidaria (e.g. Krediet et al. 2015).

Cell Density = (Cell Concentration)(Dilution )(Total Homogenate Volume) / (Sample Surface Area)

Starting database for  cell density calculations:

CytometryStarterDataset.xlsx

Example methods to get you started on cell concentration normalization:

CITATION

Koch HR, Wallace B, DeMerlis A, Clark AS, Nowicki RJ (2021). 3D Scanning as a Tool to Measure Growth Rates of Live Coral Microfragments Used for Coral Reef Restoration. Frontiers in Marine Science.

LINK

https://doi.org/10.3389/fmars.2021.623645

#### CITATION

Conley DD, Hollander ENR (2021). A Non-destructive Method to Create a Time Series of Surface Area for Coral Using 3D Photogrammetry. Frontiers in Marine Science.

LINK

https://doi.org/10.3389/fmars.2021.660846

#### CITATION

Krediet CJ, DeNofrio JC, Caruso C, Burriesci MS, Cella K, Pringle JR (2015). Rapid, Precise, and Accurate Counts of Symbiodinium Cells Using the Guava Flow Cytometer, and a Comparison to Other Methods.. PloS one.

LINK

https://doi.org/10.1371/journal.pone.0135725
